# Universal and naked-eye gene detection platform based on CRISPR/Cas12a/13a system

**DOI:** 10.1101/615724

**Authors:** Chao-Qun Yuan, Tian Tian, Jian Sun, Meng-Lu Hu, Xu-Sheng Wang, Er-Hu Xiong, Meng Cheng, Yi-Juan Bao, Wei Lin, Jie-Ming Jiang, Cheng-Wei Yang, Qian Chen, Heng Wang, Xi-Ran Wang, Xian-Bo Deng, Xiao-Ping Liao, Ya-Hong Liu, Gui-Hong Zhang, Xiao-Ming Zhou

**Author notes:** These authors contributed equally (Chao-Qun Yuan, Tian Tian, Jian Sun).

## Abstract

Colorimetric gene detection based on gold nanoparticles (AuNPs) is an attractive detection format due to its simplicity. Here, we report a new design for a colorimetric gene-sensing platform based on the CRISPR/Cas system that has improved specificity, sensitivity, and universality. CRISPR/Cas12a and CRISPR/Cas13a have two distinct catalytic activities and are used for specific target gene recognition. Programmable recognition of DNA by Cas12a/crRNA and RNA by Cas13a/crRNA with a complementary sequence activates the nonspecific *trans*-ssDNA or -RNA cleavage, respectively, thus degrading the ssDNA or RNA linkers which are designed as a hybridization template for the AuNP-DNA probe pair. Target-induced *trans* -ssDNA or RNA cleavage leads to a distance-dependent color change for the AuNP-DNA probe pair. In this platform, naked eye detection of transgenic rice, African swine fever virus (ASFV), and a miRNA can be completed within 1 hour. Our colorimetric gene-sensing method shows superior characteristics, such as probe universality, isothermal reaction conditions, on-site detection capability, and sensitivity that is comparable to that of the fluorescent detection; thus, this method represents a robust next generation gene detection platform.

---

Developing simple, inexpensive, and accurate analytical methods to detect ultralow concentrations of DNA/RNA sequences with high specificity is important for medical molecular diagnosis, agricultural and food testing, and forensic analysis(1). Ideal diagnostic tests should deliver results quickly and be enabled for instant use on a variety of sample types without excessive reliance on a technician or an auxiliary device(2, 3). Colorimetric detection using gold nanoparticles (AuNPs) is a simple and inexpensive detection format that has attracted considerable stable interest in the scientific community because aggregation of AuNPs can be clearly identified with the naked eye(4-6). Aggregation of AuNPs shifts the absorption peak to longer wavelengths and changes the color of a colloidal solution from red to purple(7). Mirkin and colleagues are the pioneers in the development of colorimetric detection strategies based on AuNPs(7, 8). They were the first to invent the DNA-labeled gold nanostructure, which they later termed spherical nucleic acids, for sequence-specific gene detection(9, 10). In the following decades, the academic community has developed a large number of colorimetric gene detection methods based on the crosslinking or non-crosslinking behavior of AuNPs(11-24).

Some of these methods are coupled with gene amplification technology; thus, the sensitivity is significantly improved and genomic samples can be analyzed. Despite great success, some of the following problems still require to be partially or fully addressed. (i) Most of the existing methods detect a red-to-purple color change, which is similar to a signal-off format. When a low concentration target is detected, the faint color change makes it difficult to detect, and the signal-to-noise ratio is very low. (ii) AuNP-DNA probes are not universal. Probes need to be redesigned when detecting different targets, thus increasing the cost and time spent for optimization of the experimental parameters. Some methods have probe universality by designing a universal tag sequence in the primer for subsequent AuNP-DNA probe hybridization(21); however, this approach causes the false positive problems due to primer dimer amplification. (iii) Detecting genomic amplification products requires denaturation of double-stranded DNA for hybridization(20). This high-temperature hybridization process is incompatible with the AuNP-DNA probe stability. Improvement of AuNP-DNA probe stability by time-consuming modification of nanoparticle surface, such as silicidation, is usually necessary(22). The problems can be addressed by employing asymmetric amplification to obtain single-stranded products for room temperature hybridization(11). However, asymmetric amplification reduces the amplification efficiency thus affecting the detection sensitivity.

Bacteria have evolved the CRISPR-Cas immune system specifically targeting exogenous threats, such as a phage or exogenous genetic material(25, 26). The sequence-specific recognition capabilities of the CRISPR system have been applied in a growing variety of fields, such as the pharmaceutical industry, the food industry, the agricultural industry and industrial biotechnology(27-29); the systems are largely developed on the basis of Cas9(30-32). Recently discovered Cas12a, Cas13a, and Cas14 system have certain new features that are different from Cas9(33-40). When Cas12a/crRNA, Cas13a/crRNA, or Cas14/crRNA recognizes their DNA or RNA, they switch to the active state, wherein they cut the single-stranded DNA/RNA substrates in a nonspecific way(33, 34, 36, 39). This collateral cleavage activity is termed *trans*-cleavage. New gene detection technologies using the CRISPR/Cas system, such as DETECTR(33), HOLMES(35), and SHERLOCK(38), with extremely high sensitivity (single molecule levels) and specificity (single base resolution) have been developed in combination with gene amplification including PCR and RPA. However, most of the strategies involve expensive double-labeled (fluorescent and quenching groups) single-stranded RNA or DNA substrates and rely on fluorescent detection equipment(33, 35, 36, 38-40). Zhang et al. developed a test strip method for onsite detection(37). However, similar to the fluorescence detection method, this method requires an expensive double-labeled substrate probe.

In an attempt to overcome these drawbacks, we developed a novel genetic detection platform based on the CRISPR technology and AuNP-DNA probes. This method has several advantages, such as probe universality, no need for expensive dual-labeled probes, isothermal reaction conditions, low cost and naked eye detection. This developed gene detection platform has inherently high specificity assisted by CRISPR recognition and maintains the simplicity of colorimetric detection. Moreover, signal-on detection mode is achieved with the assistance of a low speed centrifugation method. Successful detection of transgenic DNA, African swine fever virus (ASFV), and miRNA with sensitivity comparable to that of fluorescence methods indicates that our method is a robust next generation gene detection platform.

## Results and Discussion

### Assay Design

Recent studies have shown that Cas12a has both cis and trans single-stranded DNA cleavage activity(33, 34). When Cas12a forms a ternary complex with crRNA and its target DNA, the complex acquires a strong “chaotic” activity and cleaves single-stranded DNA into 2-4 nucleotide fragments (called trans cleavage)(34). Studies have shown that both cis-and trans-cleavage activities of Cas12a are associated with its RuvC pockets(33). Prior studies have shown that Cas13a has additional collateral cleavage activity after cleavage of its target RNA(36). Structural studies have analyzed the binary complexes of Cas13a and crRNA in Leptotrichia shahii (Lsh) bacteria, revealing that Cas13a functions through two independent active domains(41). Using this target-dependent *trans*-cleavage by Cas12a and Cas13a, we have developed a new gene detection platform based on distance-dependent optical properties of the AuNP-DNA probes (Scheme 1). In this detection platform, universal ssDNA or ssRNA linkers were designed as the *trans*-cleavage substrates for the Cas12a or Cas13a system. Additionally, a pair of universal AuNP-DNA probes was designed to hybridize to linker ssDNA or ssRNA. In the absence of a target, trans-cleavage is not activated, so the linker ssDNA and ssRNA remain intact in the reaction system. The AuNP-DNA probe pair will undergo a hybridization-induced crosslinking to form an aggregated state. When Cas12a/crRNA and Cas13a/crRNA recognize their target DNA and RNA, respectively, *trans*-cleavage is activated and the linker ssDNA and ssRNA will be degraded (Figure S1-S4). AuNP-DNA pair loses the template for hybridization and thus becomes dispersed. Visual detection can be performed by estimating the distance-dependent optical properties of the crosslinked and dispersed AuNP-DNA probes.

**Scheme 1.**
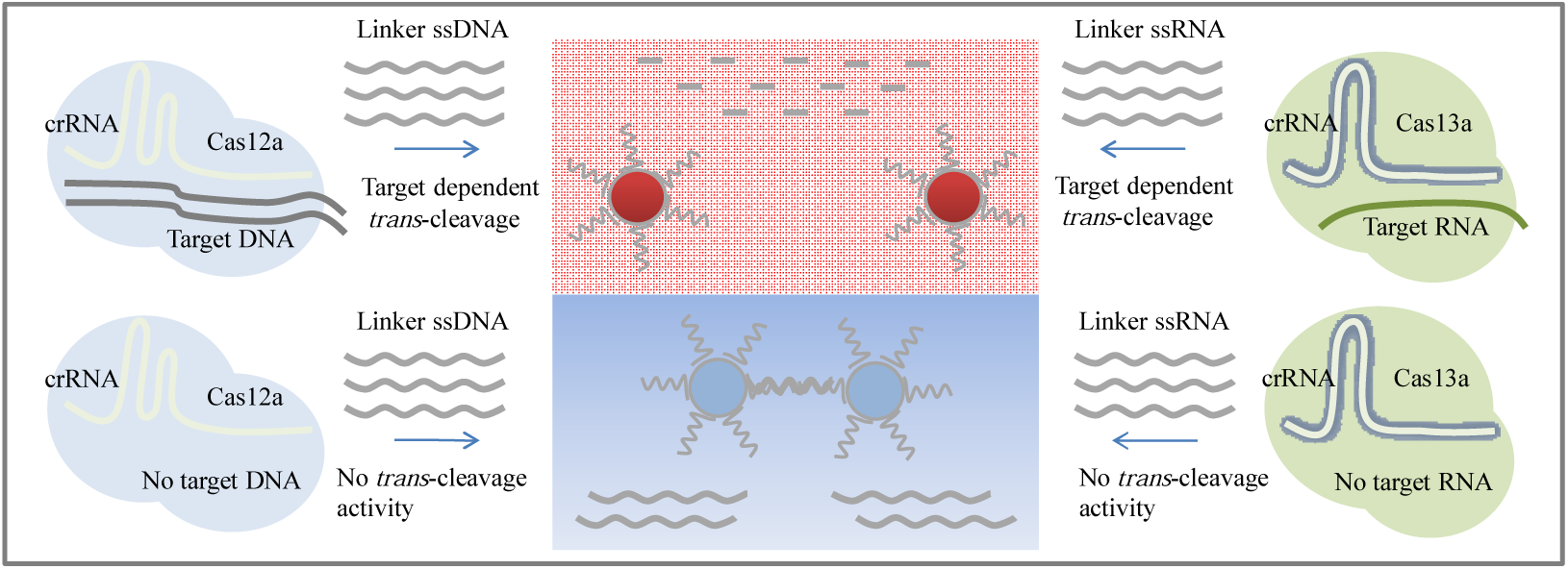
Principle scheme of CRISPR-based colorimetric gene detection. Cas12a/crRNA and Cas13a/crRNA possess target-dependent collateral cleavage activity. The collateral cleavage activity was used for signal reporting using distance-dependent optical properties of AuNP-DNA probes pair. In the presence of a target, linker ssDNA and linker ssRNA are degraded. AuNP-DNA probe pair loses the hybridization template and becomes dispersed. In the absence of a target, linker ssDNA and linker ssRNA are intact. A crosslinking reaction of the AuNP-DNA pair with linker ssDNA or ssRNA results in aggregation.

### Assay Development

Conventionally, labeling of AuNP-DNA probes is achieved by covalent conjugation of the Au-S bond(42). However, this method is time consuming, as it usually it takes tens of hours to complete a labeling reaction. Moreover, the modification of the thiol groups of the DNA probe increases the cost of the reagent. Recently reported strategy based on the affinity between poly (A) and AuNPs enables SH-modification-free AuNP-DNA labeling(43, 44). However, in this strategy, successful AuNP-DNA labeling still takes tens of hours to complete. In the current study, we show that AuNP-DNA labeling based on poly (A)-Au affinity can be accelerated by freezing and can thus be completed in two hours(45) (Figure S5). After obtaining the AuNP-DNA probe pair (Figure S6), we used hybridization experiments to evaluate the distance-dependent optical properties. However, after the AuNP-DNA probe pair was hybridized to the linker ssDNA, we did not detect a sharp red-to-blue color change regardless of the concentration of the ssDNA linker. Hybridization only causes a red-to-magenta color change. Such faint color difference makes naked eye detection difficult especially when detecting low concentrations of the targets. In principle, when the interparticle distance in these aggregates becomes less than the particle diameter, the color will become blue(19). Based on the head-to-tail hybridization pattern used, it is estimated that the distance between the AuNP-DNAs pair should be less than 13 nm. However, given that the AuNP-DNA probes used in this study were obtained by poly (A) labeling instead of Au-S labeling, we hypothesize that this surface-covering labeling method increases the actual interparticle distance between the AuNP-DNA pair after hybridization. It is shown that this hybridization reaction resulted in the formation of a network-like crosslinked complex with a size change from approximately 20 nm to about 700 nm (Figure S7). The crosslinked AuNP-DNA probes and the AuNP-DNA monomer can be easily separated by a low-speed centrifugation step. Experiments demonstrated that only 3 minutes of low speed centrifugation (5,000 rpm) are required to complete the precipitation of crosslinked AuNP-DNA probes resulting in the apparently colorless and transparent supernatant of the probe solution (Figure S8). The uncrosslinked AuNP-DNA probe solution remained in a red colloid state (Figure 1A). Therefore, the low speed centrifugation method allows the achievement of an easily distinguishable color change and a high signal-to-noise ratio. Considering that low speed centrifuges are very easy to obtain and inexpensive (approximately $100) (Figure S8), this method greatly enhances the colorimetric detection ability without sacrificing the detection convenience.

**Figure 1.**
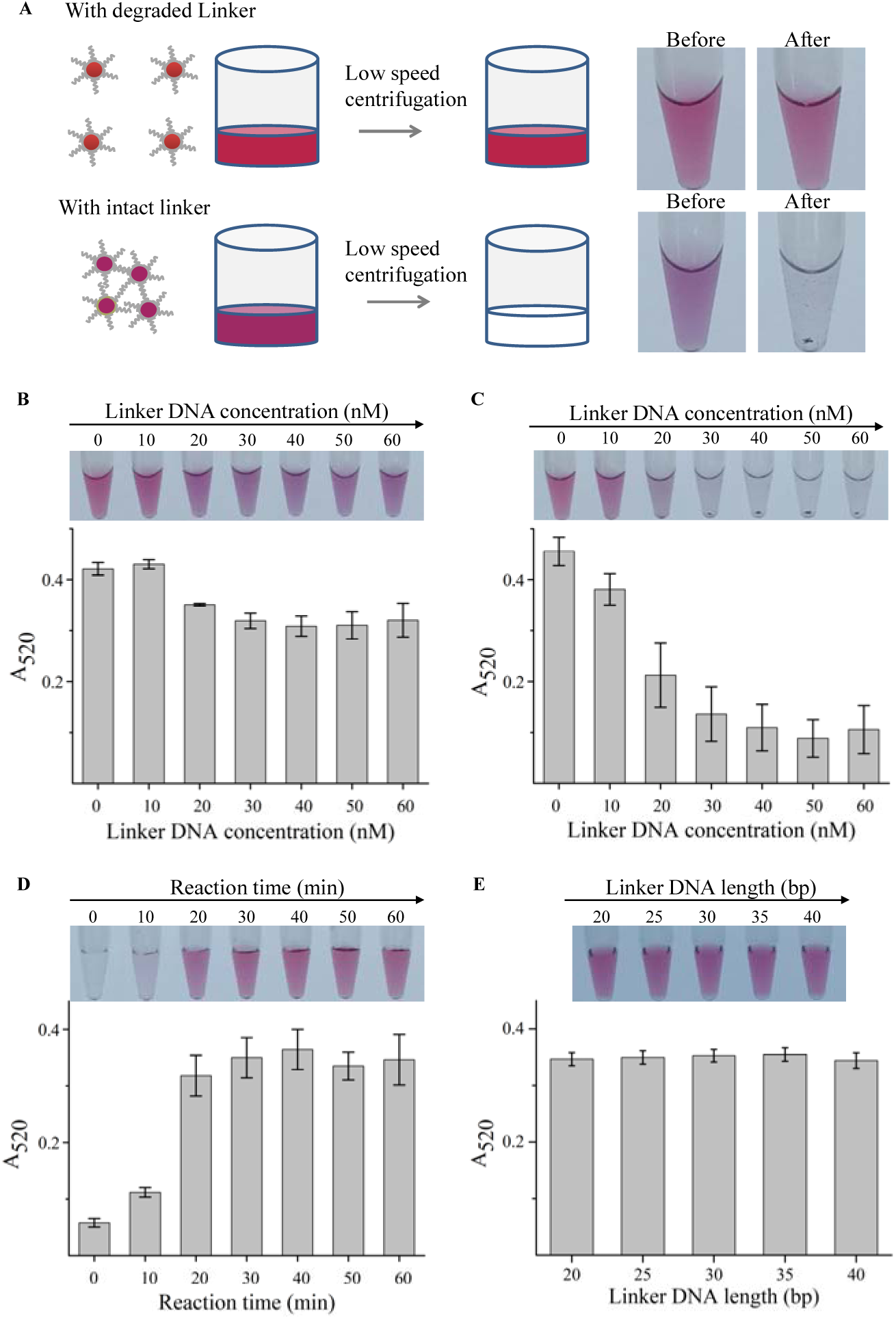
Assay development of CRISPR-based colorimetric gene detection. (A) Low speed centrifugation-assisted colorimetric detection. Due to degradation of the linker DNA, the AuNP-DNA probe pairs loses the hybridization template thus becoming highly dispersive. Color does not change before and after low speed centrifugation. With intact linker DNA, hybrid crosslinking between the AuNP-DNA probe pair and linker DNA causes a color change from red to magenta. Low speed centrifugation of the crosslinked complexes deposits them at the bottom of the tube, thus, the solution becomes colorless and transparent. (B) Screening of the critical linker DNA concentration by direct observation of color change and measurement of absorption at 520 nm. (C) Screening of the critical linker DNA concentration by low speed centrifugation and measurement of absorption at 520 nm. (D) Monitoring of reaction dynamics of the trans-cleavage in the Cas12a/crRNA system by AuNP-DNA probe pair with the low speed centrifugation method. (E) Monitoring of linker DNA length dependence of the Cas12a/crRNA system by AuNP-DNA probe pair with the low speed centrifugation method.

Next, we decided to determine the linker DNA concentration that would be sufficient to cause the full crosslinking of the AuNP-DNA probes. Target-dependent *trans*-cleavage capability of the Cas system is positively correlated with the target DNA/RNA concentration; hence, linker DNA/RNA substrate concentration will be the key factor determining the detection sensitivity. Using linker DNA as an example, we tested the crosslinking behavior of the AuNP-DNA pair (5 nM) at 0, 10, 20, 30, 40, 50, and 60 nM linker DNA concentrations. The critical linker DNA concentration for complete AuNP-DNA crosslinking was evaluated by direct observation of color changes and low speed centrifugation followed with a measurement of the absorption of AuNP-DNA probe solution at 520 nm. As shown in Figure 1B and 1C, direct observation and the low speed centrifugation methods indicate that the crosslinking reaction of the AuNP-DNA pair is saturated at linker DNA concentrations of greater than 30 nM. However, very narrow observation window is detected in the case of direct observation. With low speed centrifugation, a very sharp color change can be obtained. At linker DNA concentration of 40 nM, a sufficiently clean background signal can be obtained, and the signal-to-noise ratio is significantly improved. Then, using the EGFP gene as a target, we evaluated the time kinetics of target-dependent trans-cleavage activity of Cas12a/crRNA in the presence of 40 nM linker. It was found that linker DNA cleavage can be completed in 20 minutes (Figure 1D). This indicates that Cas12a/crRNA-based detection system requires only 20 minutes of incubation time. Finally, we tested another parameter, the length of the linker ssDNA, which may be an important factor for determining the sensitivity. A previous study has demonstrated that Cas14 has a functional similarity to Cas12a and showed strong preference for cleavage of the longer substrates(39). In the current experiments, we evaluated linker ssDNA of various lengths (20, 25, 30, 35, and 40 bp) and found that Cas12a does not have this length dependency (Figure 1E). Considering that longer sequences may form secondary structures that affect hybridization at room temperature, we selected a 20 bp linker ssDNA for subsequent use.

### Workflow for CRISPR-based colorimetric gene detection

After the method was developed, we proceeded to design a workflow for the gene detection platform (Scheme 2). Because Cas/crRNA complexes, linker DNA or RNA substrate and AuNP-DNA probes can be prepared in advance and are suitable for long-term storage, the whole detection process can be summarized in three steps: preparation of solution 1, preparation of solution 2, and colorimetric detection. The preparation of solution 1 includes mixing the target genes with the Cas/crRNA complex and linker substrate; trans-cleavage reaction of the Cas system is activated by the target genes, and in total, the procedure takes approximately 20 minutes. The preparation of solution 2 is very simple and requires simple mixing of the AuNP-DNA probe pair. This process takes less than 2 minutes and can be performed while waiting for preparation of solution 1. Finally, a drop of solution 1 is added to solution 2 and after a 3-minute incubation followed by low speed centrifugation, the test results can be observed with the naked eye. Colorimetric detection takes less than 30 minutes. In combination with the RPA technology, genome sample analysis can be completed within an hour. Nucleic acid samples with various numbers of copies can be determined by observing the grades of the color of the AuNP-DNA probe.

**Scheme 2.**
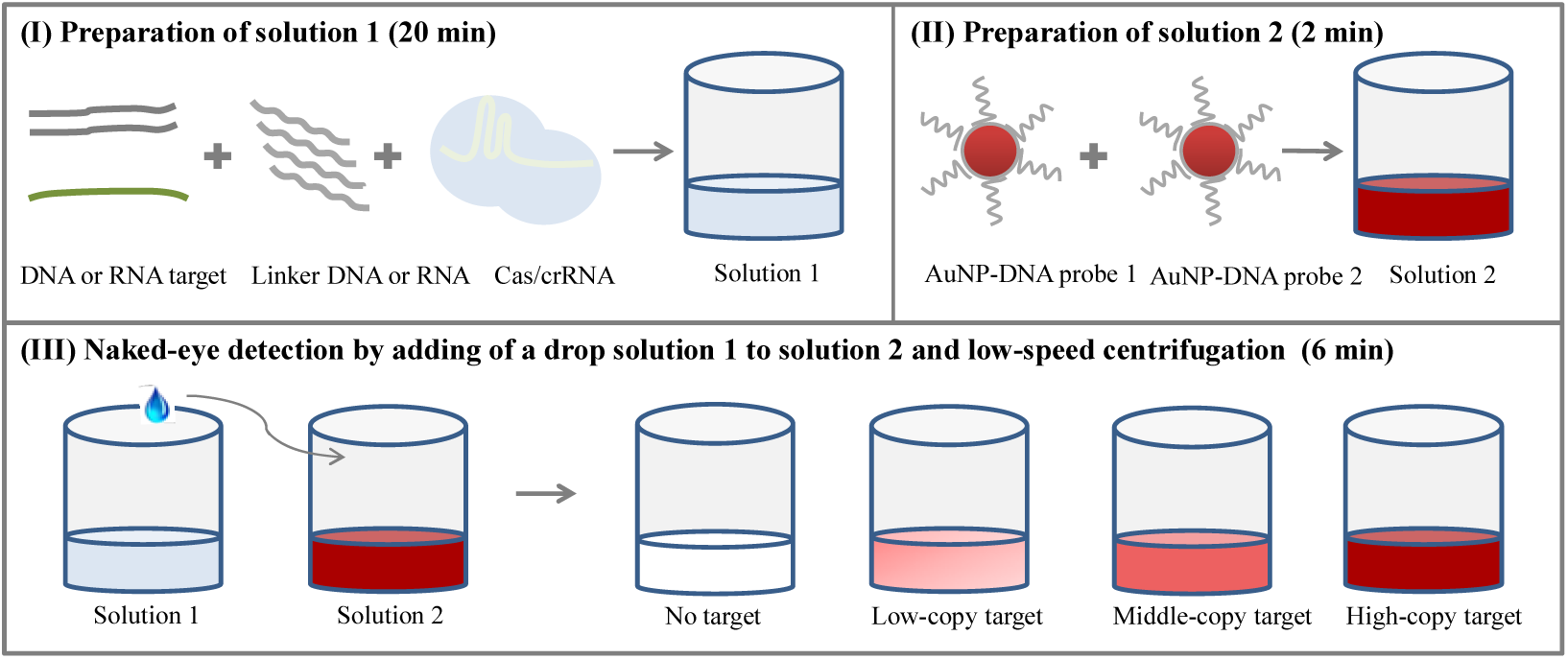
Workflow of CRISPR-based colorimetric gene detection. (A) Target DNA and RNA are added to a Cas/crRNA complexes in the presence of linker ssDNA or ssRNA and incubated for 20 min to prepare solution 1. (B) AuNP-DNA probes pair is mixed to prepare solution 2. (C) Naked eye detection can be completed by adding of a drop of solution 1 to solution 2 followed by low speed centrifugation.

### CRISPR-based colorimetric gene detection of GMOs

To strengthen the management of genetically modified crops, more than 40 countries and regions including the European Union have enacted the corresponding laws and regulations for management of genetically modified organisms (GMOs) and their products(46). For example, the EU requires a mandatory labeling system when the genetically modified ingredients exceed 1%. The most common components of the transgenic construct, the promoter of the cauliflower mosaic virus (p35S) and the nopaline synthase gene from Agrobacterium tumefaciens (NOS), has become the typical targets for GMO screening methods(47). This study is the first to apply the CRISPR-based colorimetric detection system to the detection of 35S promoter from transgenic rice. Figure 2A shows that Cas12a/crRNA recognizes a 20 bp target sequence in a 35S sequence adjacent to a TTTA PAM site. *Trans*-cleavage by Cas12a/crRNA is activated by recognition of the 35S sequence with the detection of the signal output using the AuNP probe. Figure 2B shows that straightforward GMO detection is achieved by a combination with PCR or RPA amplification. Compared to electrophoretic analysis, the results of CRISPR-based colorimetric detection are easier to evaluate. Compared to the reported colorimetric detection methods that usually involve a change of color from red to reddish-purple to blue, the current method is based on the detection of a colorless–to-red change thus representing a signal-on detection format. Moreover, CRISPR-based colorimetric detection involves a second step of probe recognition (primer pair amplification represents the first step of recognition), so it can exclude nonspecific signals from the primer dimers or target-independent amplification.

**Figure 2.**
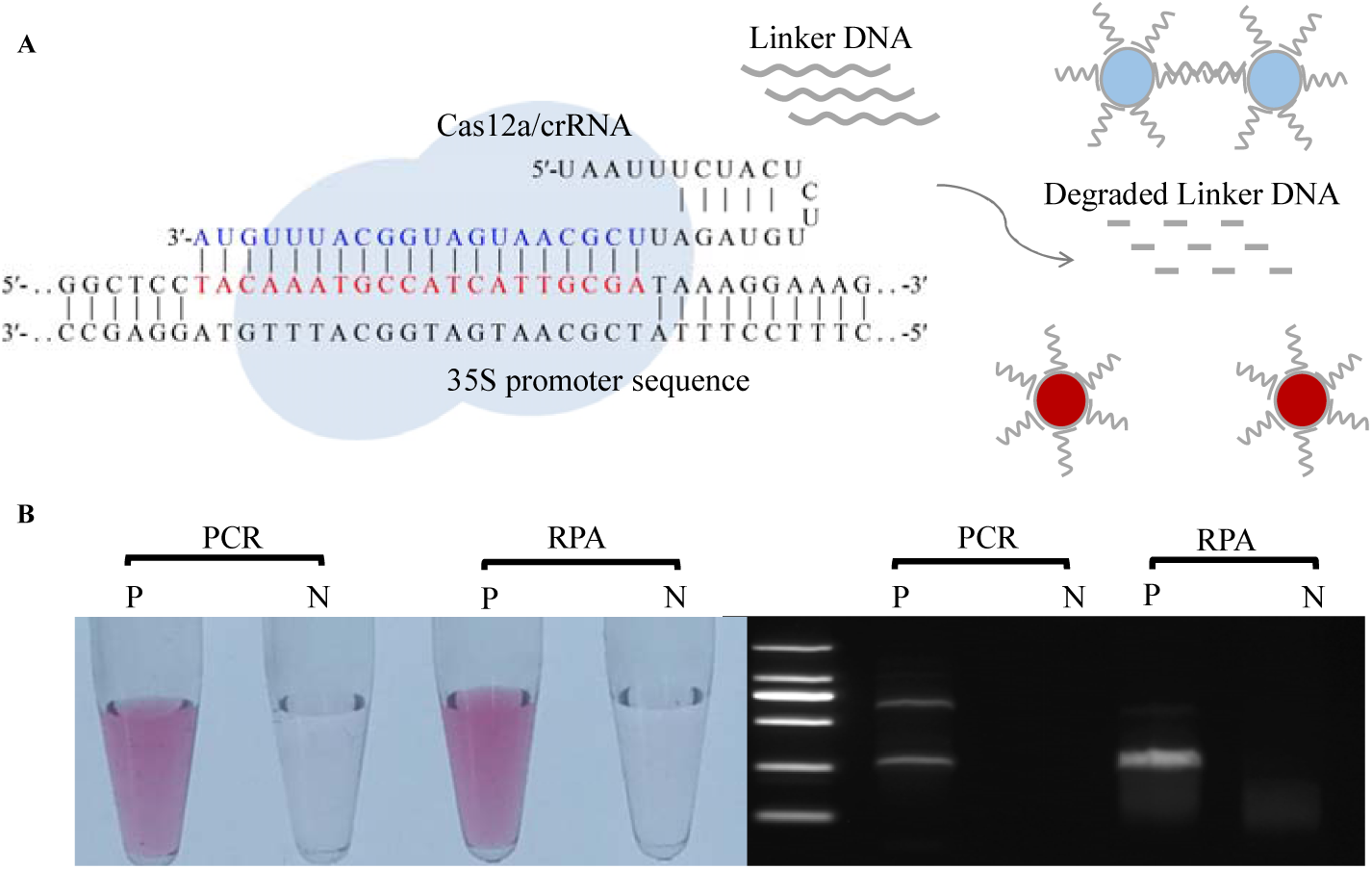
CRISPR-based colorimetric gene detection of GMOs. (A) 35S promoter sequence recognition by the Cas12a/crRNA system followed by colorimetric analysis. (B) Colorimetric analysis of the RPA and PCR products of the 35S promoter (left). Electrophoretic analysis of the RPA and PCR products of the 35S promoter (right). P and N stands for positive and negative samples, respectively.

Next, we evaluated the detection ability of the CRISPR-based colorimetric detection system for 35S. We mixed transgenic rice with non-transgenic rice and milled the rice into powder for nucleic acid extraction. Combined with PCR and RPA, CRISPR-based colorimetric assays can detect transgenes in rice at as low as the 0.01% level (Figure 3C and 3D). The sensitivity of colorimetric detection is one or two orders of magnitude higher than that of the electrophoresis analysis (Figure 3A and 3B). The results show that the sensitivity of the method is fully qualified for the needs of the required detection of transgenic GMO components ranging from 0.1 to 1% worldwide.

**Figure 3.**
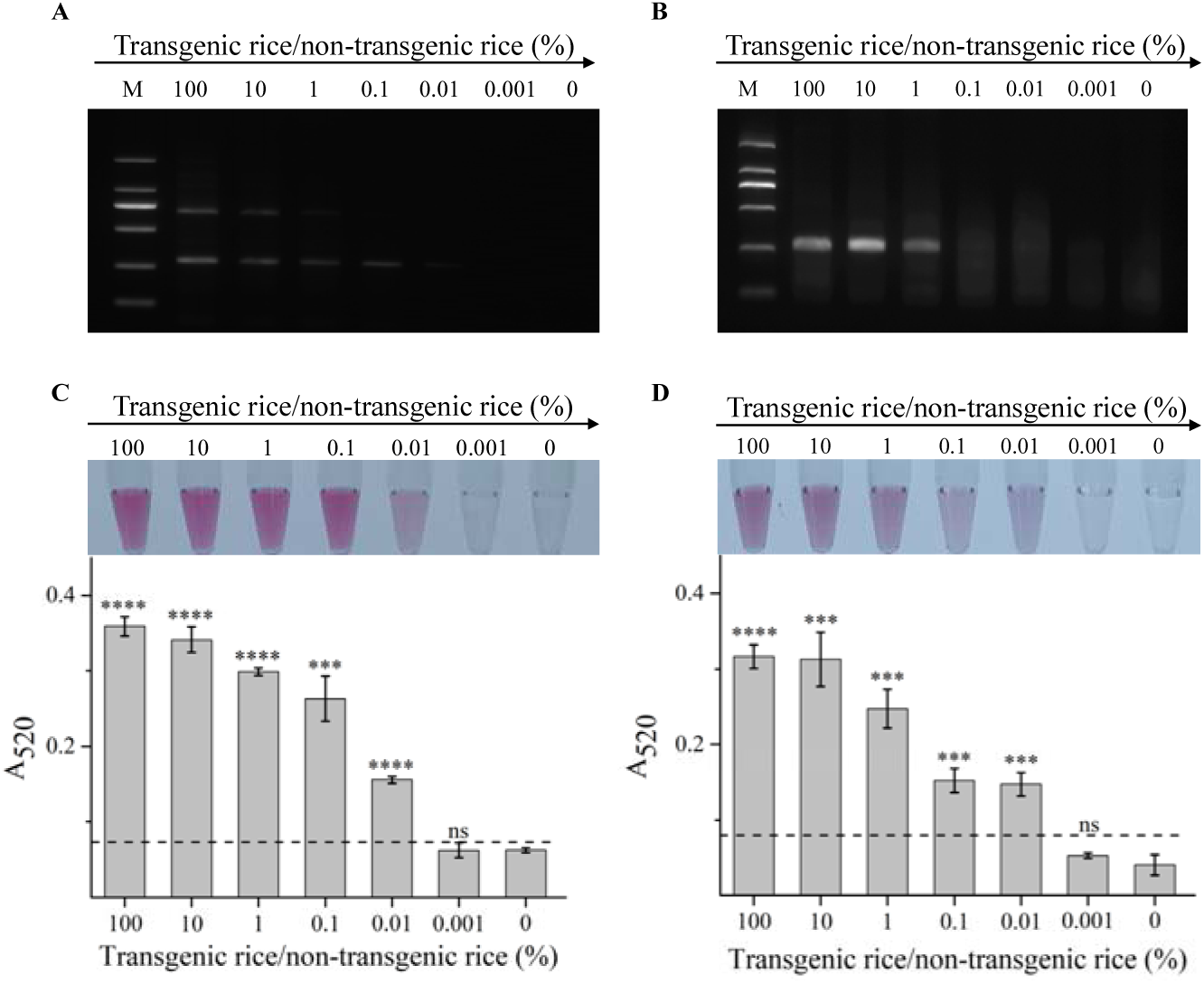
Electrophoretic and CRISPR-based colorimetric analysis of PCR and RPA products from different percentage of transgenic rice DNA. (A) Electrophoretic analysis of RPA products with varied percentage of transgenic rice DNA. (B) Electrophoretic analysis of PCR products with varied percentage of transgenic rice DNA. (C) Colorimetric analysis of RPA products. Photograph showing colorimetric responses of the detection system in the presence of different percentage of transgenic rice DNA. The picture below represents the absorbance (A520) signals of the colorimetric analysis. (D) Colorimetric analysis of the PCR products. Photograph showing colorimetric responses of the detection system in the presence of different percentage of transgenic rice DNA. The picture below represents the absorbance (A520) signals of the colorimetric analysis. The threshold line represents the mean ± three standard deviations; *p < 0.05, **p < 0.01, and ***p < 0.001.

### CRISPR-based colorimetric gene detection of ASFV

African swine fever (ASF) is an acute and potent hemorrhagic infectious disease of the pigs caused by African swine fever virus (ASFV)(48). ASFV is a double-stranded DNA virus with a capsular membrane belonging to the African porcine virus family(49). ASFV is transmitted through feces and air and is highly contagious and has a 100% mortality rate(48, 49). Since 2018, ASF has expanded into more than 14 countries(50, 51). At present, there is no effective treatment or vaccine for ASF. Therefore, early diagnosis of ASF for prevention and control is extremely important. Rapid detection of ASFV infection can be achieved by immunological methods(52). However, these methods have a sensitivity problem because in acute infections, pigs usually die before becoming antibody-positive. This study applied the developed colorimetric gene detection to the rapid and sensitive diagnostic of ASFV to help combat the spread of the fatal disease on a global scale. We used RPA or PCR to amplify the VP72 gene of ASFV. VP72 gene encodes the p72 protein, the major capsid protein of ASFV and the main component of the viral icosahedral capsid, which is highly antigenic and immunogenic. We then designed Cas12a/crRNA to recognize the VP72 gene sequence. VP72 gene binds to the designed Cas12a/crRNA complexes to activate *trans*-cleavage resulting in disassembly of the AuNP-DNA probes and subsequent easily detectable color change (Figure 4A). Detection sensitivity was evaluated by naked eye detection or by absorption measurement, and the limit of detection was 200 copies (Figure 4B and 4C).

**Figure 4.**
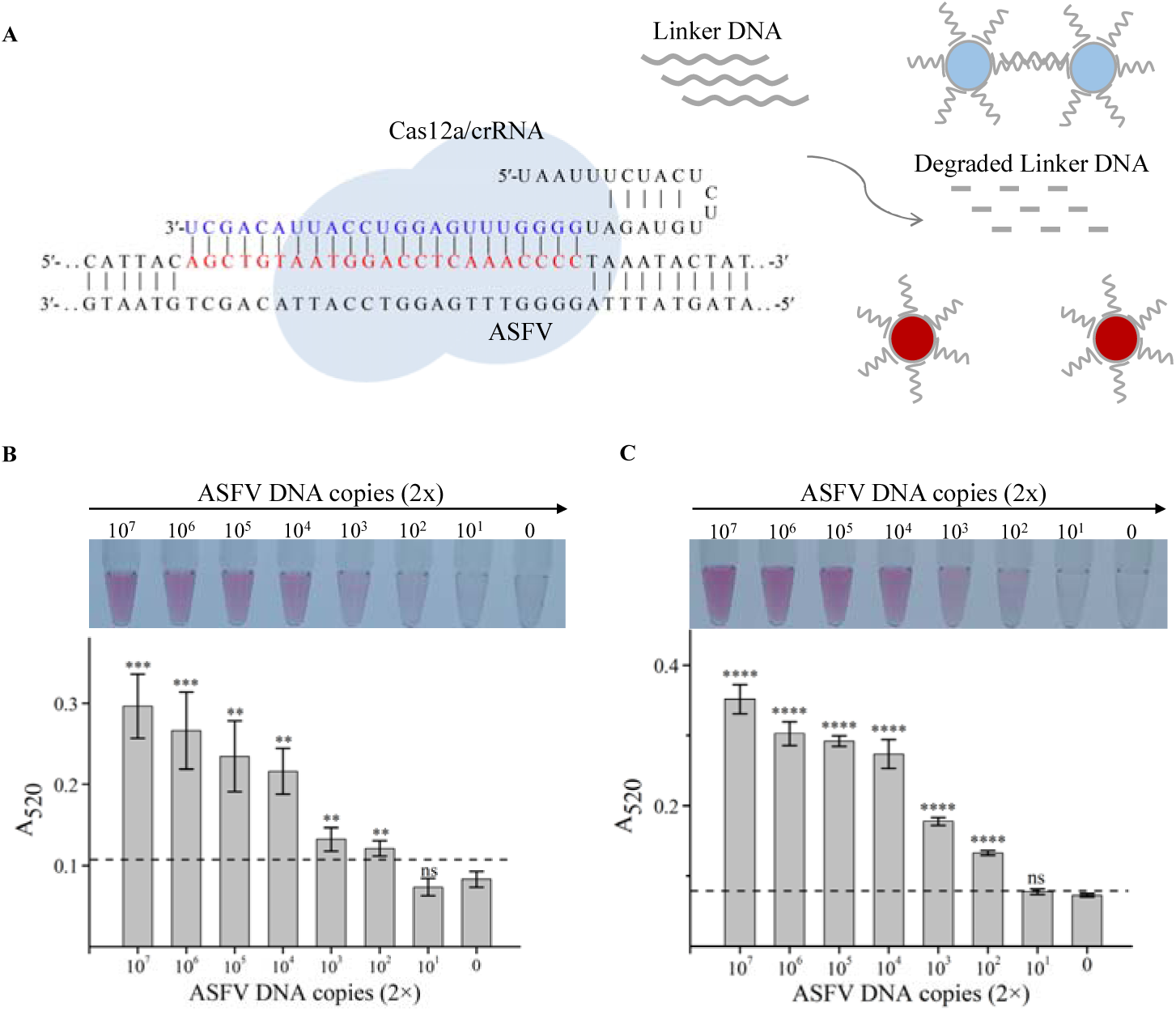
CRISPR-based colorimetric gene detection of ASFV. (A) ASFV gene sequence (VP72) recognition by the Cas12a/crRNA system and scheme of colorimetric analysis. (B) Colorimetric analysis of the PCR products. Photograph showing colorimetric responses of the detection system in the presence of different number of ASFV DNA copies. The picture below represents the absorbance (A520) signals of colorimetric analysis. (C) Colorimetric analysis of the RPA products. Photograph showing colorimetric responses of the detection system in the presence of different number of ASFV DNA copies. The picture below represents the absorbance (A520) signals of colorimetric analysis. The threshold line represents the mean ± three standard deviations; *p < 0.05, **p < 0.01, and ***p < 0.001.

### Extension to RNA detection

Cas13a/crRNA and Cas12a/crRNA have similar target-dependent *trans*-cleavage activity. Therefore, we hypothesized that the developed colorimetric detection platform can be easily extended to RNA detection by employing the same AuNPs-DNA probe pair. Cas13a/crRNA recognizes a target RNA and activates *trans*-cleavage with very high efficiency. For example, a *trans*-cleavage efficiency of at least 10^4^ turnovers per target RNA recognized was observed in the case of LbuCas13a, which is a robust enzyme with a high turnover rate(36). Our previous studies have shown that miRNAs at low fM concentrations can be detected using the dual-labeled fluorescent RNA substrates(53). This potent detection ability prompted direct miRNA detection without an amplification step. This feature is very important because miRNAs are very short and some indirect steps, such as reverse transcription and PCR, will have an adverse effect on the detection of their true expression levels.

In this study, we applied the CRISPR-based colorimetric method to RNA detection. RNA detection using this system only needs minor modifications compared with the CRISPR/Cas12a-based DNA colorimetric detection strategy. As shown in Figure 5A, using miRNA-17 as an example, the Cas13a/crRNA complex is designed to probe the miRNA17 sequence. A linker ssRNA with same sequence as an ssDNA linker used in the CRISPR/Cas12a-based colorimetric detection system, was employed as the substrate for target-dependent trans-cleavage. The sensitivity of miRNA-17 detection was as low as 500 fM (Figure 5B). This sensitivity is comparable or even superior to the methods based on dual-labeled fluorogenic probes (Figure S9). We attribute this high sensitivity to the clean background signal obtained by low speed centrifugation. Moreover, the colorimetric method had good selectivity for evaluation of target miRNA-17 compared with other interfering agents such as miRNA-10b, miRNA-17, and miRNA-155 (Figure 5C).

**Figure 5.**
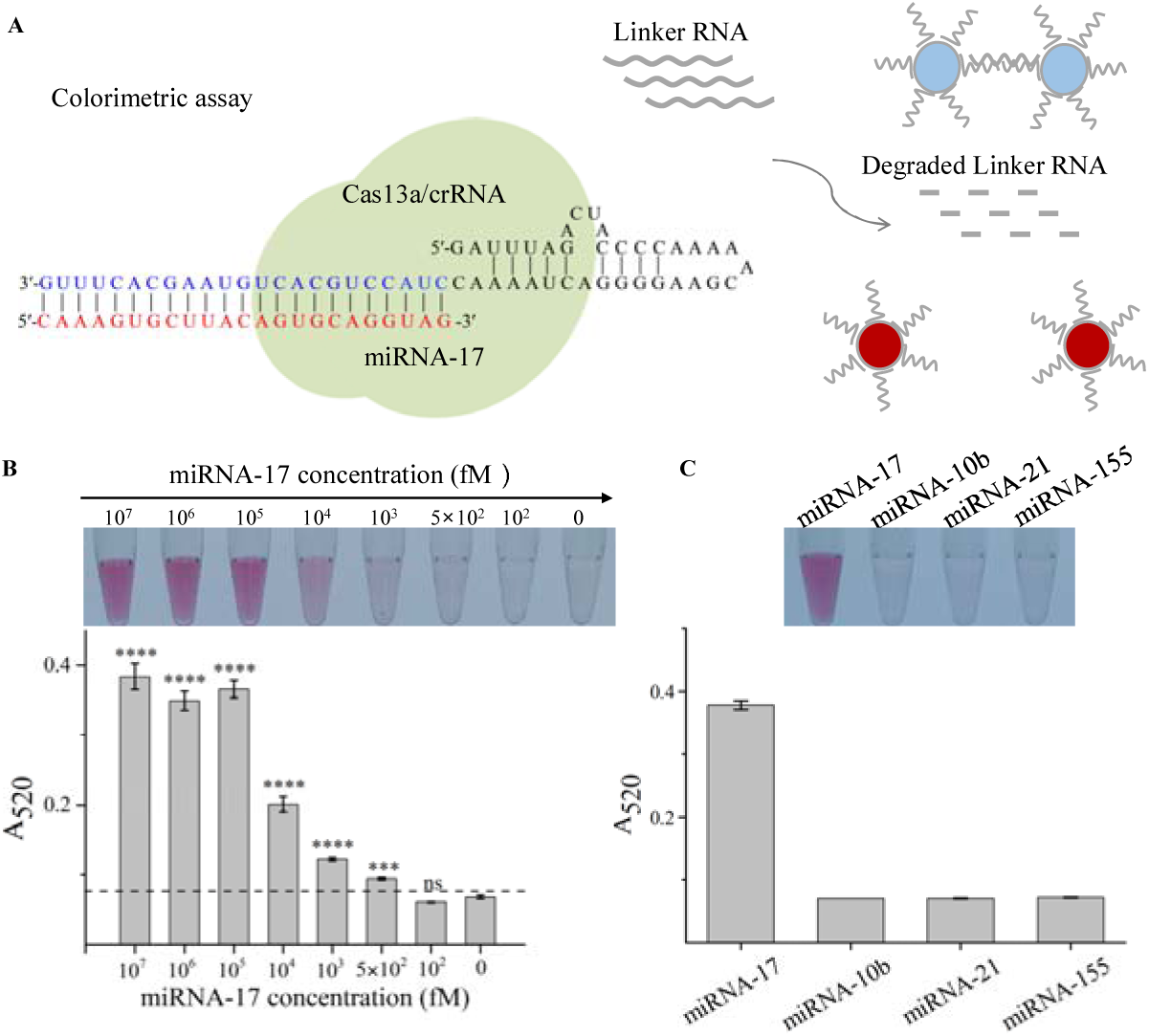
CRISPR-based colorimetric RNA detection. (A) miRNA-17 recognition by the Cas13a/crRNA system and scheme of colorimetric analysis. (B) Colorimetric analysis of miRNA-17. Photograph showing colorimetric responses of the detection system in the presence of different concentrations of miRNA-17. The picture below represents the absorbance (A520) signals of colorimetric analysis. (C) Photographs showing colorimetric detection of various target miRNA strands. The picture below represents the absorbance (A520) signals of colorimetric analysis. The threshold line represents the mean ± three standard deviations; *p < 0.05, **p < 0.01, and ***p < 0.001.

## Conclusion

Colorimetric gene detection based on gold nanoparticle has been developed for more than 20 years. However, certain disadvantages, such as signal-off detection format, low signal-to-noise ratio, requirements for high temperature hybridization, lack of probe universality, and low discrimination of color changes, are encountered during the use of the system for detection of genomic samples and hinder their applicability. Here, we presented a colorimetric gene detection platform based on the CRISPR/Cas12a/13a system that has achieved several critical advances compared to the existing method: (i) No high temperature hybridization process is required and specificity is significantly improved with the introduction of isothermal CRISPR/Cas recognition; (ii) The use of low speed centrifugation to assist in development of a signal-on colorimetric gene detection with improved detection sensitivity. Detection sensitivity of our colorimetric assay is comparable to that of the fluorescence methods. (iii) Development of thiol modification-free AuNP-DNA labeling method accelerated by freezing, which can be completed in two hours. The developed strategy represents an improvement in time and cost control. (iv) Universality of the current colorimetric gene detection platform was demonstrated by successful detection of transgenic rice, African swine fever virus, and a miRNA with the same AuNP-DNA probe. The presented gene detection platform addresses the shortcomings of existing colorimetric methods and is very promising as an onsite diagnostic tool.

## Acknowledgment

This work was supported by the National Natural Science Foundation of China (Grant 21475048; 21874049), the National Science Fund for Distinguished Young Scholars of Guangdong Province (Grant 2014A030306008), and the Special Support Program of Guangdong Province (Grant 2014TQ01R599).

