## Supporting Information for "Universal and naked-eye gene detection platform based on CRISPR/Cas12a/13a system"

a. College of Biophotonics & College of Life Science, South China Normal University, Guangzhou, Guangdong, 510531, China
b. College of Veterinary Medicine, South China Agricultural University, Guangzhou 510642, China
c. National Risk Assessment Laboratory for Antimicrobial Resistance of Animal Original Bacteria, South China Agricultural University, Guangzhou, China.
d. Research Center for African Swine Fever Prevention and Control, South China Agricultural University, Guangzhou, China.
e. Guangdong Provincial Key Laboratory of Biotechnology for Plant Development, School of Life Science, South China Normal University, Guangzhou 510631, China
f. School of Chemistry and Materials Science, Jiangsu Normal University, Xuzhou 221116, China
1. These authors contributed equally.

Experimental Procedures

**Experimental materials and instruments：**

All chemicals were from Guangzhou Chemical Reagent Factory. The enzymes and NTPs used for RNA transcription and the buffers were purchased from Bio-Lifesci (Guangzhou, China). PCR reagents were purchased from Takara (Japan). RPA reagent was obtained from TwistDx Inc. (England). All DNA sequences were synthesized by Sangon biotech (Shanghai), while the fluorescent double-labeled probe was synthesized by Takara (Japan). All DNA and RNA sequences used in this study are provided in Table S1. HAuCl_4_ solution was obtained from Aladdin (Shanghai, China). Reagents used for protein expression and purification were purchased from Abiotech (Jinan, China). The 6His-MBP-TEV-huAsCpf1 used for preparation of AS Cas12a was a gift from Feng Zhang (Addgene plasmid # 90095; http://n2t.net/addgene:90095; RRID: Addgene_90095). The p[lasmid](http://dict.youdao.com/w/plasmid/#keyfrom=E2Ctranslation) used for the expression of LbuCas13a was a gift from Professor Yanli Wang (Institute of Biophysics, Chinese Academy of Sciences). Fluorescence detection was performed on a thermal cycler Dice™ Real Time System III (Takara, Japan). Absorption spectroscopy detection was performed on a SpectraMax iD5 multi-mode microplate reader (Molecular Devices). The particle size was measured by a Zetasizer Nano-ZS instrument (Malvern Instruments). RPA experiment was performed in a thermostatic metal bath (MIULAB, Hangzhou). Low-speed centrifugation experiment was carried out using a mini benchtop centrifuge (Sangon Biotech, Shanghai). High speed centrifugation and PCR experiments were performed using the instruments from Eppendorf. Agarose and PAGE electrophoresis experiments were performed using a Beijing Liuyi instrument and imaged by a gel imager (Baijing, Beijing). All other buffers and solutions were prepared using ultrapure water (>18.25 MΩ).

**Synthesis and characterization of AuNPs:**

AuNPs were prepared by the sodium citrate reduction method as reported previously. Before the experiment, the glassware used was immersed in aqua regia (HNO_3_:HCl = 1:3) for over 30 minutes and then rinsed with ultrapure water. To a 250 ml flask, 100 mL of 1 mM HAuCl_4_ was added and heated to boiling on a heated magnetic stirrer. After boiling became stable, 10 mL of 38.8 mM sodium citrate solution was added with rapid stirring. The solution became colorless after a short time. After 20 minutes, the solution turned wine red and the heating was stopped. Then, the solution was cooled to room temperature to obtain AuNP solution (13 nm in diameter). AuNP solution had the maximum absorption at 520 nm. AuNPs were observed by electron microscopy using a TEM (JEM-2100HR) to determine the particle size. The prepared AuNPs solution was stored at 4 °C in the dark until further use.

**Labeling of AuNP-DNA probes by freezing:**

To 1 mL of AuNPs solution, 50 μL of 100 μM poly(A)-tagged DNA probe was added. The solution was mixed and frozen in a refrigerator (-20 °C). The freezing time was at least 2 hours. After thawing, the solution was centrifuged at 4 °C at 12,000 rpm for 30 minutes. The supernatant was aspirated and the precipitate was resuspended in a wash buffer (0.1 M NaCl in 0.01 M phosphate buffer, pH 7.4). This step was repeated three times. Finally, the pellet was resuspended in a buffer (0.3 M NaCl in 0.01 M phosphate buffer, pH 7.4) and the AuNP-DNA probe was stored at 4 °C in the dark.

**Expression and purification of Cas12a and LbuCas13a proteins:**

AsCas12a and LbuCas13a proteins were expressed in E. coli Rosetta 2 (DE3) cells and cultured overnight in the TB medium containing chloramphenicol and ampicillin. Then, IPTG was added and induction was performed at 37 °C for 4 hours. The collected bacterial solution was centrifuged to obtain a precipitate and lysed by sonication in 20 mM Tris-HCl, pH 7.5, containing 1 M NaCl, 20 mM imidazole, and 10% glycerol. After centrifugation at 4 °C, the supernatant was purified using a nickel column (Abiotech, Jinan, China) and eluted with eluent 1 (20 mM Tris-HCl, pH 7.5, 150 mM NaCl, and 250 mM imidazole). The eluted AsCas12a protein was digested with TEV protease to remove the 6His-MBP tag and then purified again using a heparin column (Abiotech, Jinan, China), and the protein was eluted with eluent 2 (20 mM Tris-HCl, pH 7.5, 1 M NaCl, and 10% glycerol). Protein was collected and concentrated; then, glycerol was added to the final concentration of 50%. The protein was stored at -20 °C for further use.

**Preparation of crRNA**

crRNA was obtained by in vitro transcription. The DNA templates were heated to 95 °C and then slowly cooled to room temperature for annealing. In a 50 μL transcription system, NTPs (final concentration of 0.5 mM), 250 U T7 RNA polymerase, 50 U recombination RNase inhibitor and 200 ng of template DNA were added and incubated for 4 hours at 37 °C. Then, DNase I was added to digest the excess DNA template; the transcripts were purified by an RNA purification kit and stored at -80 °C for further use.

**Fluorescence detection of target DNA based on Cas12a**

In a 20 μL reaction system, Cas12a reaction was carried out at a final concentration of 100 nM AsCas12a, 200 nM crRNA, 200 nM DNA fluorescent probe, and varied concentrations of the DNA amplification products. Fluorescence signals were recorded by a thermal cycler DiceTM Real Time System III. The reaction was carried out at 37 °C for 30 minutes and the fluorescence signal was recorded every minute.

**Fluorescence detection of target RNA based on Cas13a**

In a 20 μL reaction system, Cas13a reaction was carried out at a final concentration of 100 nM LbuCas13a, 200 nM crRNA, 200 nM RNA fluorescent probe, and varied concentrations of the target RNA. Fluorescence signals were recorded by a thermal cycler DiceTM Real Time System III. The reaction was carried out at 37 °C for 30 minutes, and the fluorescence signal was recorded every minute.

**Rice genome extraction:**

The transgenic and nontransgenic rice samples were pulverized into powder in a mortar. Genomic DNA extraction was performed using a plant genomic DNA kit. When extracting different proportions of transgenic rice DNA, the transgenic and nontransgenic rice powders were prepared according to different mass ratios (for example, 10% transgenic rice sample is prepared by mixing 1 g transgenic rice powder and 9 g nontransgenic rice powder). The mixed rice powder was used for genome extraction. The extracted genomic samples were stored at -20 °C until use.

**Preparation of nucleic acids of ASFV:**

Viral DNA templates were provided by Guangdong Provincial Centers for Disease Control and Prevention of China. These clinical samples were used as a source of the viral DNA templates and were collected from Huangpu District, Guangzhou City, Guangdong Province.

**Amplification of 35S promoter of transgenic rice by PCR or RPA:**

PCR amplification of the 35S promoter was performed using a TaKaRa Ex Taq® kit. For example, a 20 μL amplification system contained a primer pair at a final concentration of 300 nM and varied concentrations of the genome samples. The PCR conditions were: 95 °C for 10 min followed by 35 cycles of amplification reaction at 95 °C for 30 s, 55°C for 30 s, and 72°C for 40 s with a final extension at 72°C for 10 min. The PCR product was then used for colorimetric detection or for agarose gel electrophoresis analysis. The RPA amplification of the 35S promoter was performed using a TwistAmp kit. For example, a 20 μL amplification system contained a primer pair at a final concentration of 300 nM and varied concentrations of the genome samples. The RPA reaction conditions were 37°C for 20 min. The RPA product was then used for colorimetric detection or for agarose gel electrophoresis analysis.

**Amplification of VP72 gene of ASFV by PCR or RPA:**

PCR amplification of the VP72 gene of ASFV was performed using a TaKaRa Ex Taq® kit. For example, a 20 μL amplification system contained a primer pair at a final concentration of 300 nM and varied concentrations of the genome samples. The PCR conditions were: 95 °C for 10 min followed by 35 cycles of amplification reaction at 95 °C for 15 s, 55 °C for 30 s, and 72 °C for 45 s with a final extension at 72°C for 10 min. The PCR product was then used for colorimetric detection or for agarose gel electrophoresis analysis. The RPA amplification of the VP72 gene of ASFV was performed using a TwistAmp kit. For example, a 20 μL amplification system contained a primer pair at a final concentration of 420 nM and varied concentrations of the genome samples. The RPA reaction conditions were 37 °C for 20 min. The RPA product was then used for colorimetric detection or for agarose gel electrophoresis analysis.

**Colorimetric DNA detection based on CRISPR/Cas12a:**

Cas12a/crRNA reaction buffer: 100 mM NaCl, 50 mM Tris-HCl, 10 mM MgCl_2_, and 100 μg/ml BSA, pH 7.9. A 20 μL reaction system contained 100 nM AsCas12a, 200 nM crRNA, and 200 nM linker DNA substrate. DNA amplification product (2 μL) was added to the solution, mixed and reacted in a metal bath for 15 minutes at 37°C; then, 60 μL of solution was prepared using equal volumes of the two premixed AuNP-DNA probe solutions (30 μL of AuNP-DNA 1 and 30 μL of AuNP-DNA 2); the samples were incubated at room temperature for 3 min and centrifuged in a tabletop centrifuge at 5,000 rpm for 3 min. The supernatant was used for colorimetric detection over a white backlight plate.

**Colorimetric RNA detection based on CRISPR/Cas13a:**

Cas13a/crRNA reaction buffer: 100 mM NaCl, 50 mM Tris-HCl, 10 mM MgCl_2_, and 100 μg/ml BSA, pH 7.9. A 20 μL reaction system contained 100 nM LbuCas13a, 200 nM crRNA, and 200 nM linker RNA substrate. Target RNA (2 μL) was added to the solution, mixed and reacted in a metal bath for 15 minutes at 37°C; then, 60 μL of solution was prepared by adding equal volumes of two premixed AuNP-DNA probe solutions (30 μL of AuNP-DNA 1 and 30 μL of AuNP-DNA 2); the samples were incubated at room temperature for 3 min and centrifuged in a tabletop centrifuge at 5,000 rpm for 3 min. The supernatant was used for colorimetric detection over a white backlight plate.

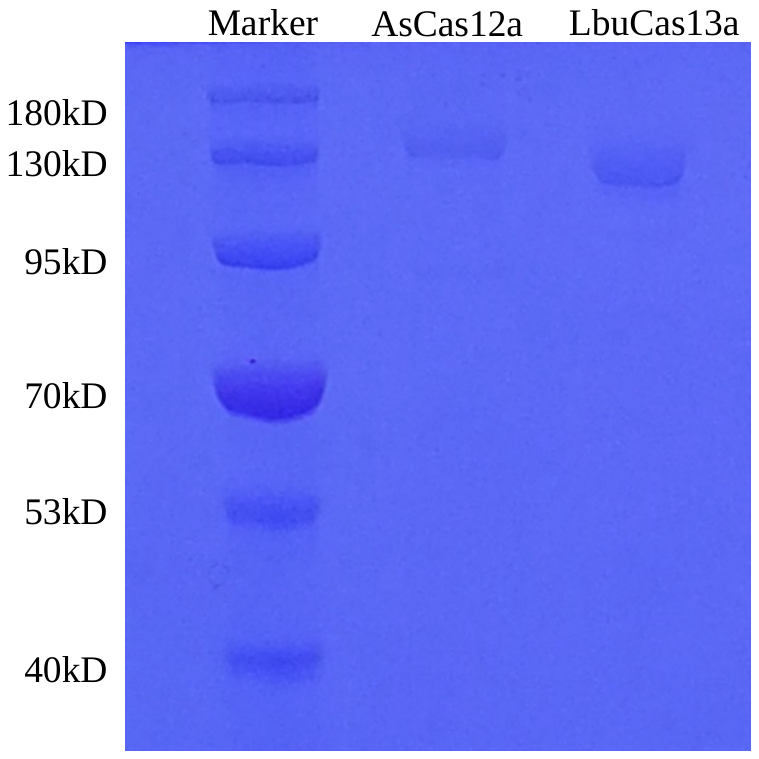

Figure S1. PAGE gel analysis of AsCas12a and LbuCas13a. Two dilutions of AsCas12a and LbuCas13a are shown.

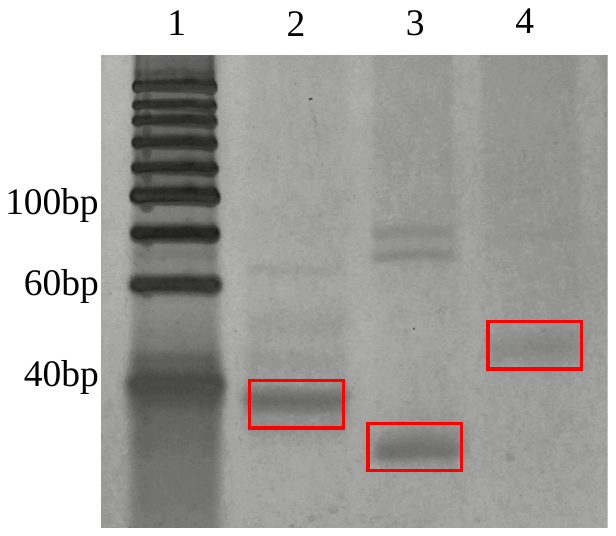

Figure S2. PAGE gel analysis of synthesized crRNAs. Line 2, 3, and 4 represent crRNA for targeting 35S promoter, ASFV, and miRNA-17, respectively.

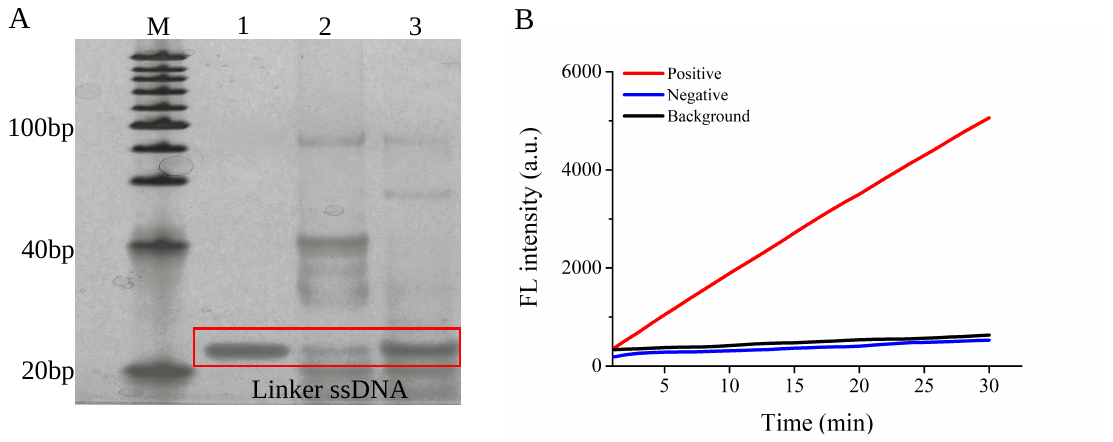

Figure S3. Characterization of *trans*-cleavage of Cas12a system. (A) PAGE gel analysis of *trans*-cleavage of Cas12a system. Line1: linker ssRNA, line 2: Cas12a/crRNA+ target DNA+ linker ssDNA; line 3: Cas12a/crRNA+ linker ssDNA; (B). Fluorescence analysis of *trans*-cleavage of Cas12a system. Positive: Cas12a/crRNA+ target DNA+ ssDNA Taqman probe; Negative: Cas12a/crRNA+ ssDNA Taqman probe; Background: ssDNA Taqman probe.

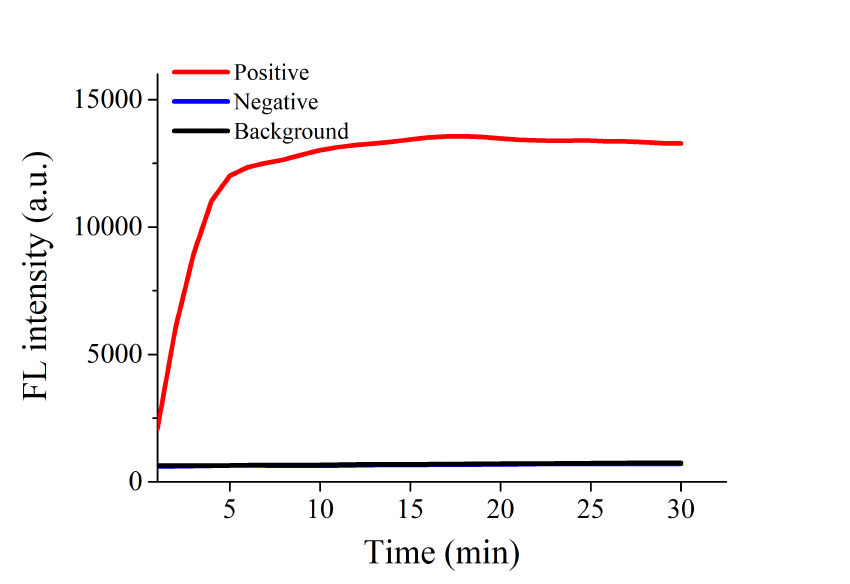

Figure S4. Characterization of *trans*-cleavage of Cas13a system. Fluorescence analysis of *trans*-cleavage of Cas13a system. Positive: Cas13a/crRNA+ target RNA+ ssRNA Taqman probe; Negative: Cas13a/crRNA+ ssRNA Taqman probe; Background: ssRNA Taqman probe.

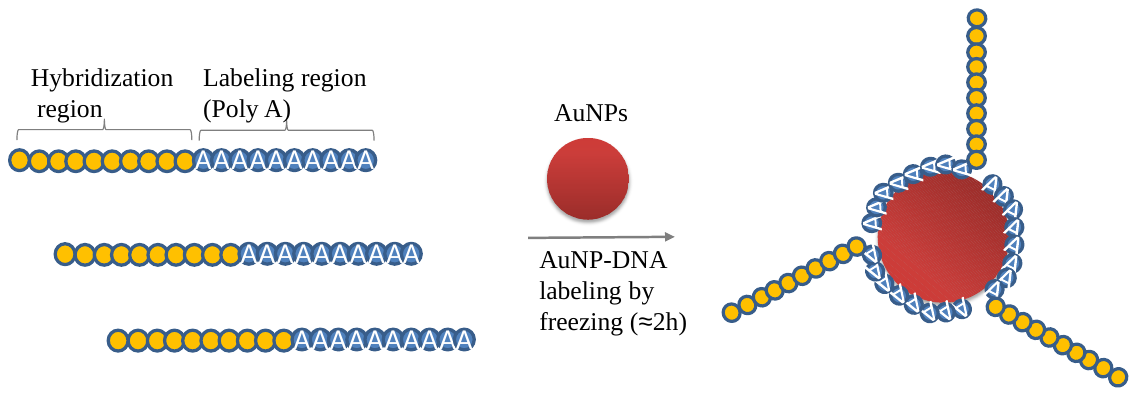

Figure S5. AuNP-DNA labeling based on poly (A)-Au affinity accelerated by freezing. The process can be completed in 2 hours.

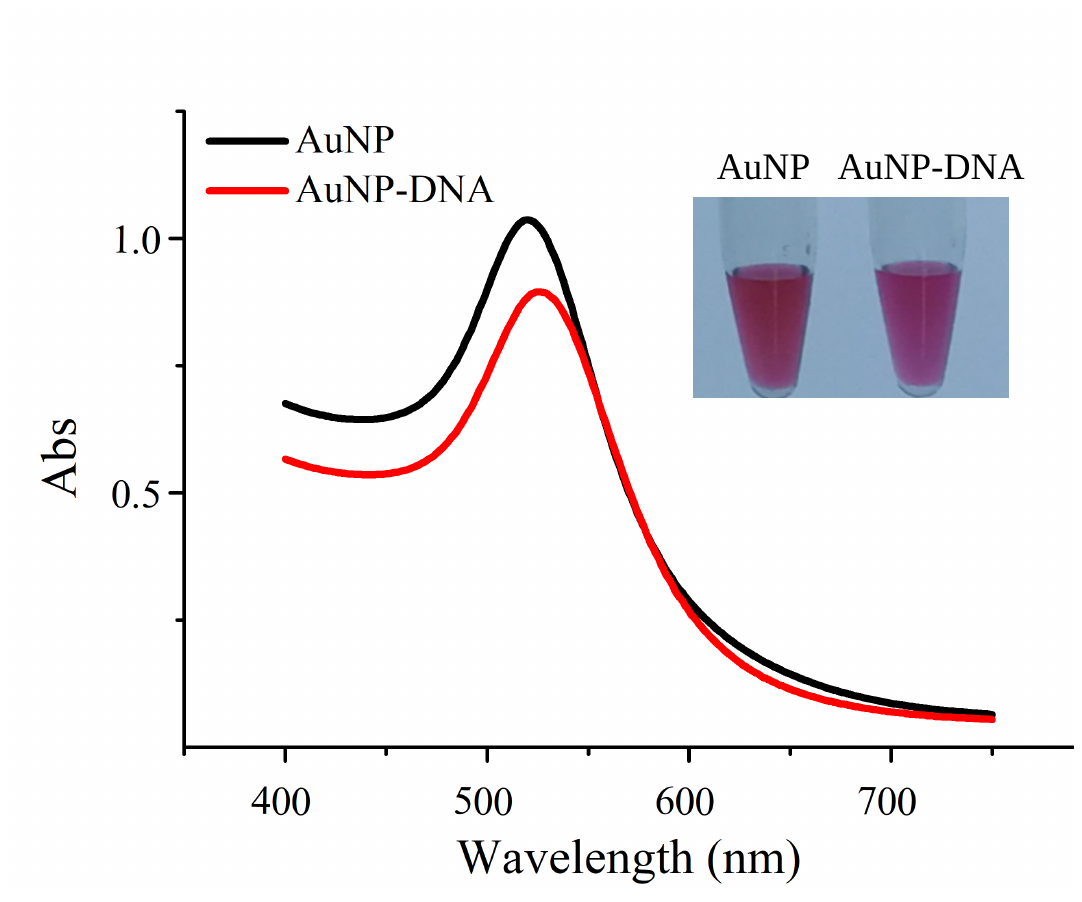

Figure S6. Characterization of AuNP-DNA labeling with a measurement of the absorption. [Upper right](http://dict.youdao.com/w/upper%20right/#keyfrom=E2Ctranslation) shows a [picture](http://dict.youdao.com/w/picture/#keyfrom=E2Ctranslation) before and after DNA labeling.

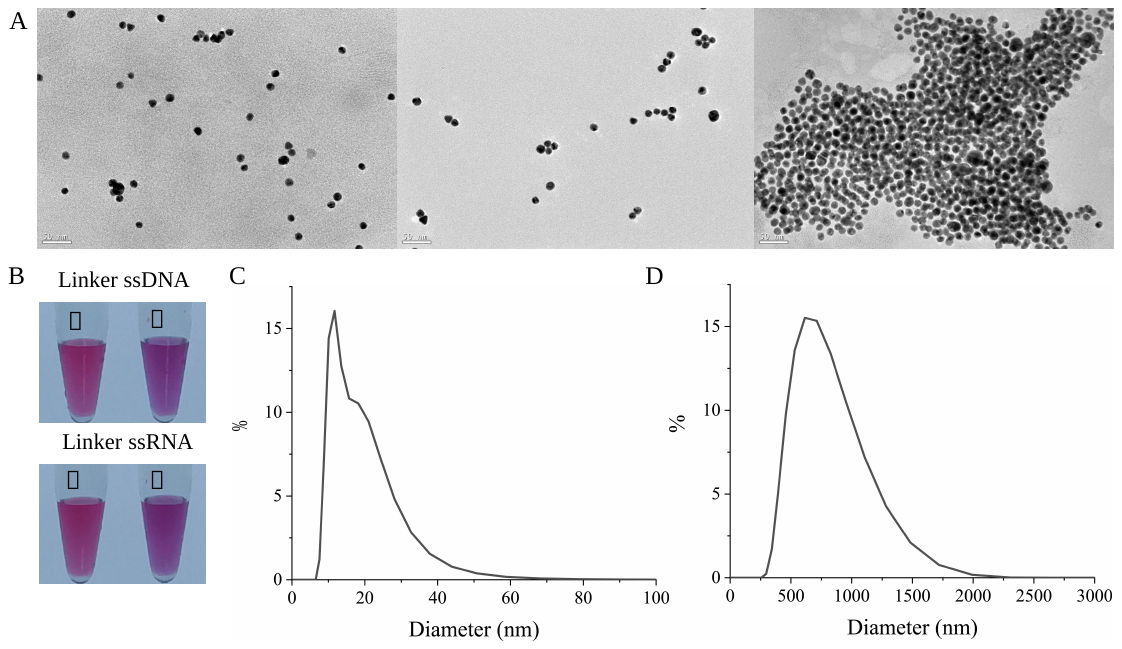

Figure S7. Characterization of linker DNA or RNA caused crosslink of AuNP-DNA probe. (A) [Electron microscope](http://dict.youdao.com/w/electron%20microscope/#keyfrom=E2Ctranslation) analysis of naked AuNPs (left), AuNPs-DNA probe (middle), and crosslinked AuNP-DNA probe (right); (B) [Picture](http://dict.youdao.com/w/picture/#keyfrom=E2Ctranslation)s show the color change of AuNP-DNA probe pair before and after crosslinked by linker ssDNA and ssRNA. (C) Size analysis of AuNPs-DNA probe by [dynamic light scattering](http://dict.youdao.com/w/dynamic%20light%20scattering/#keyfrom=E2Ctranslation). (D) Size analysis of crosslinked AuNPs-DNA probe by [dynamic light scattering](http://dict.youdao.com/w/dynamic%20light%20scattering/#keyfrom=E2Ctranslation).

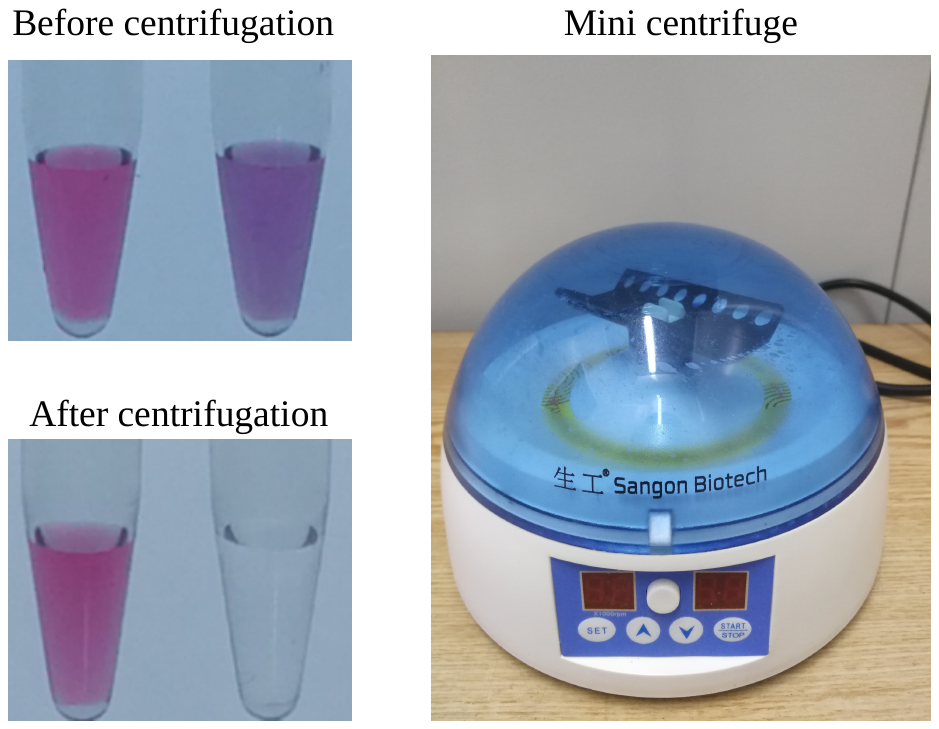

Figure S8. Crosslinked and uncrosslinked AuNP-DNA probe pair solution observed by naked eye. Before centrifugation the solution shows a red-to-[purplish red](http://dict.youdao.com/w/purplish%20red/#keyfrom=E2Ctranslation) color change. After centrifugation the solution shows a red-to-colorless change. Right shows the centrifuges used in this method.

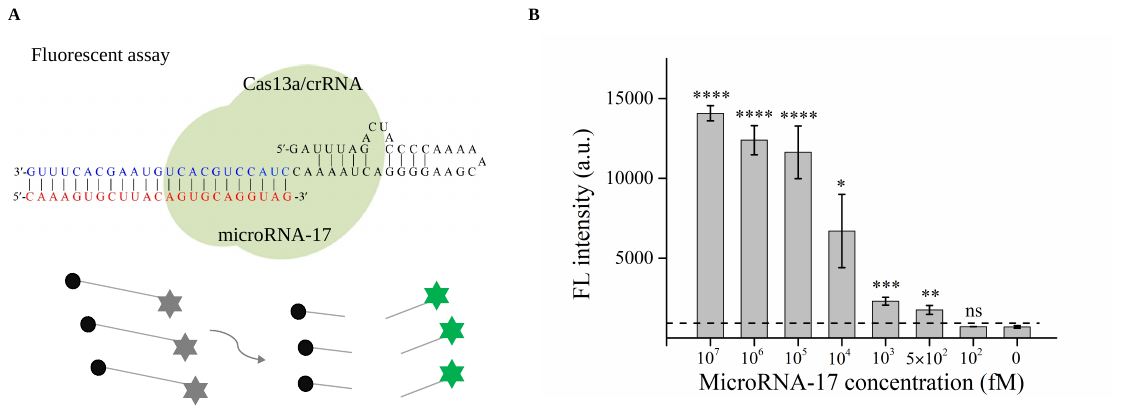

Figure S9. (A) Scheme of fluorescent analysis of miRNA-17 based on the Cas13a/crRNA system. (B) Statistical results of fluorescence signal measured at different concentrations of miRNA-17. *p < 0.05, **p < 0.01, and ***p < 0.001.

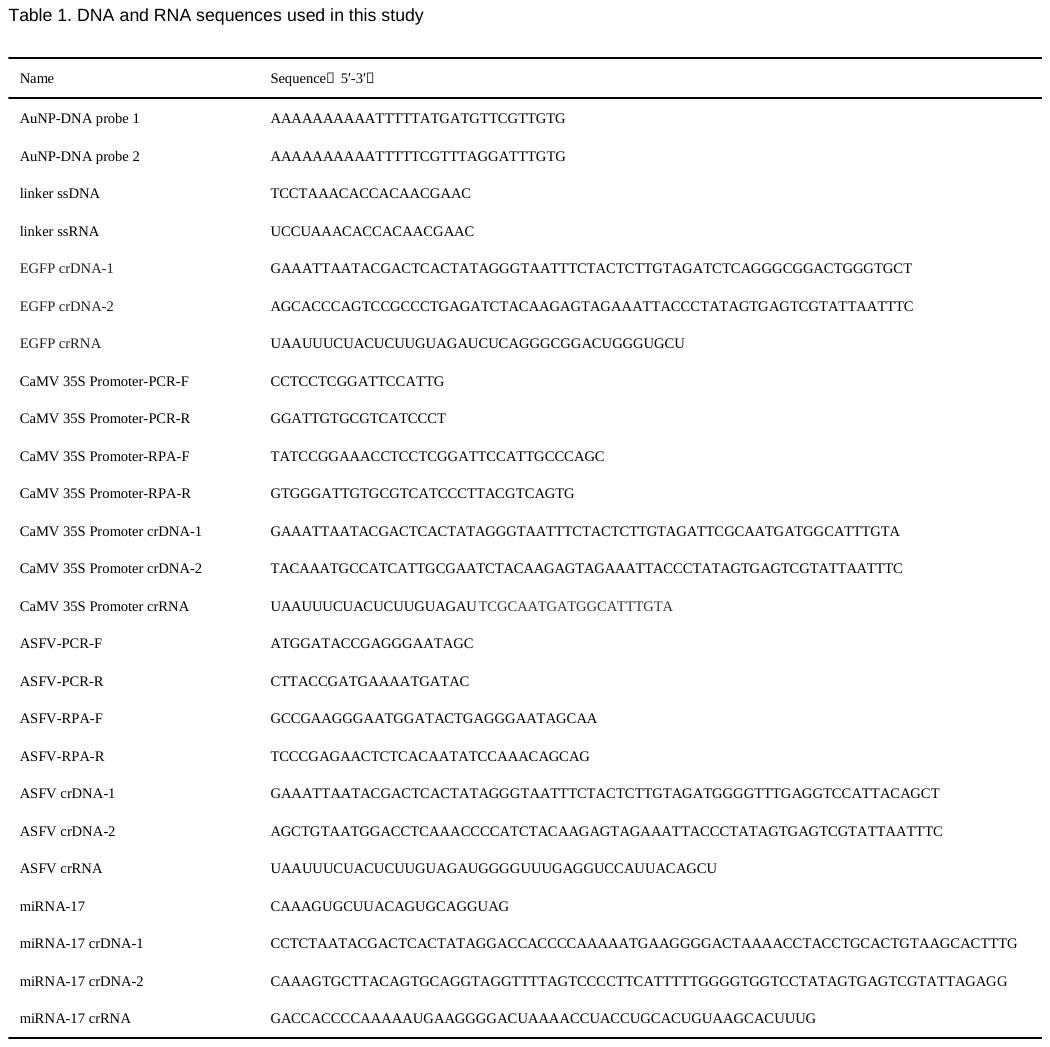
